# Protein family annotation for the Unified Human Gastrointestinal Proteome by DPCfam clustering

**DOI:** 10.1101/2023.04.21.537802

**Authors:** Federico Barone, Elena Tea Russo, Edith Natalia Villegas Garcia, Marco Punta, Stefano Cozzini, Alessio Ansuini, Alberto Cazzaniga

## Abstract

Technological advances in massively parallel sequencing have led to an exponential growth in the number of known protein sequences. Much of this growth originates from metagenomic projects producing new sequences from environmental and clinical samples. The Unified Human Gastrointestinal Proteome (UHGP) catalogue is one of the most relevant metagenomic datasets with applications ranging from medicine to biology. However, the lack of sequence annotation impairs its usability. This work aims to produce a family classification of UHGP sequences to facilitate downstream structural and functional annotation. This is achieved through the release of the *DPCfam-UHGP50 dataset* containing 10,778 putative protein families generated using DPCfam clustering, an unsupervised pipeline grouping sequences into multi-domain architectures. DPCfam-UHGP50 considerably improves family coverage at protein and residue levels compared to the manually curated repository Pfam. It is our hope that DPCfam-UHGP50 will foster future discoveries in the field of metagenomics of the human gut by the release of a FAIR-compliant database easily accessible via a searchable web server and Zenodo repository.

## Background & Summary

In recent years, advances in sequencing technology and the consequent drop in cost and time of experiments resulted in an exponential growth of the number of known protein sequences. For example, the latest release of the UniProtKB database, containing 227M sequences, expands the 2019 version by 90% [30, 7]. Metagenomic databases grow even faster: the EMBL-EBI MGnify, including hundreds of millions of sequences produced from high-quality assembled genomes, reported a 48-fold growth since its first release in 2017 [24][19].

Functional annotation of protein sequences is crucial for a broad spectrum of biological applications, ranging from understanding cellular mechanisms to drug discovery [14, 25]. Since large-scale experimental characterization is unfeasible, researchers rely on alternative tools to impute sequence-function relations. A prominent strategy consists of grouping homologous protein regions into families to formulate functional hypotheses based on common ancestry. Current efforts to provide protein family annotations rely at least to some degree on manual intervention, thus limiting their ability to match the ever-increasing influx of sequenced data while inherently introducing human bias. For example, the number of families in the Pfam database, focused on classifying all protein domains in evolutionary-related groups, increased by 6% over the last five years [8, 16]. Despite the introduction of novel families, Pfam maintained a steady level of sequence and residue coverage of the UniProtKB database over the same time period, respectively 77% and 53%. Increasing the level of coverage with the inclusion of novel families is becoming more challenging as new Pfam entries tend to cover a small taxonomic range [8, 16]. Furthermore, the proliferation of metagenomic projects will likely induce a further drop in Pfam coverage. In fact, family annotation levels of metagenomic protein sequence databases are low, typically around 46% [24].

To address this problem, we developed the DPCfam pipeline, a two-step Density Peak Clustering (DPC) algorithm [26] which classifies sequence-similar protein regions into putative protein families called metaclusters (MCs) [28]. Crucially, the pipeline generates the final classification through a fully unsupervised algorithm that relies only on local pairwise sequence alignments. Supported by a high-performance modular implementation that ensures scaling over large databases, DPCfam consists of a fully automated alternative tool that can improve the annotation of protein datasets. In previous work, we successfully used DPCfam to classify the UniRef50 dataset (version 2017_07), containing about 23 million sequences[27]. As part of this work, 45,000 MCs were generated, of which 30% consisted of protein regions that lacked Pfam annotation. These regions may represent novel protein families. The results from this procedure have already been used to identify 63 new protein families in Pfam 35.0.

In the present study, we apply the DPCfam pipeline to metagenomics data, focusing on the Unified Human Gastrointestinal Protein catalogue clustered at 50% amino acid identity (UHGP-50). This dataset is part of the MGnify catalogue and contains ∼4 million entries representing ∼600 million proteins encoded in the Unified Human Gastrointestinal Genome (UHGG) collection [1]. The overall scope of the Unified Human Gastrointestinal initiative was to generate a nonredundant dataset of the human gut genomes and proteins by merging together the most relevant studies [10, 6, 32, 9, 33] into a unified curated catalogue. The dataset has been widely used by the scientific community and has already fostered important applications in medicine and biology [29, 22]; despite this, large portions of the catalogue lack functional annotation.

Running the DPCfam pipeline on UHGP-50 generated 10,778 metaclusters, each consisting of a set of sequence-similar protein regions called seeds. We filter seed regions based on sequence similarity for each MC to build profile-HMMs [17]. These profiles were then used to perform a more sensitive search on UHGP-50. We took all significant (domain E-value=0.03, protein E-value=0.01) profile-HMM hits as our new MC member regions. Using this extended MC representation, DPCfam covers 50.02% of all residues and 53.38% of all sequences in UHGP-50. This represents a substantial gain in coverage if compared with the Pfam annotation reported in the original UHGP release [1], which has a residue and sequence coverage for UHGP-50 of 26.2% and 37.55% respectively. Of all metaclusters, 1,261 do not overlap either with families in Pfam or with metaclusters in DPCfam-UniRef50 and thus represent potential novel families, which might be gut metagenome-specific.

The aim of this study is to provide researchers with a database that could accelerate the identification of novel protein families in the Human Gastrointestinal Proteome, thus potentially enabling the formulation of new functional hypotheses. Future plans include further optimization of the clustering procedure allowing for faster processing of newly deposited metagenomic data, and combining automatic classification with functional annotation leveraging deep learning [3, 31] for even more comprehensive annotation of UHGP.

## Methods

We present a schematic overview of DPCfam in Fig. 1. The pipeline consists of four stages: 1) all-versus-all sequence alignment; 2) domain identification, also referred to as first clustering; 3) family creation, also referred to as secondary clustering or metaclustering; 4) metacluster merging and filtering procedure. Pre- and post-processing steps are needed depending on each specific case. We briefly review each stage of the pipeline, and we refer to the work of Russo et al. [27] for further details.

**Figure 1:**
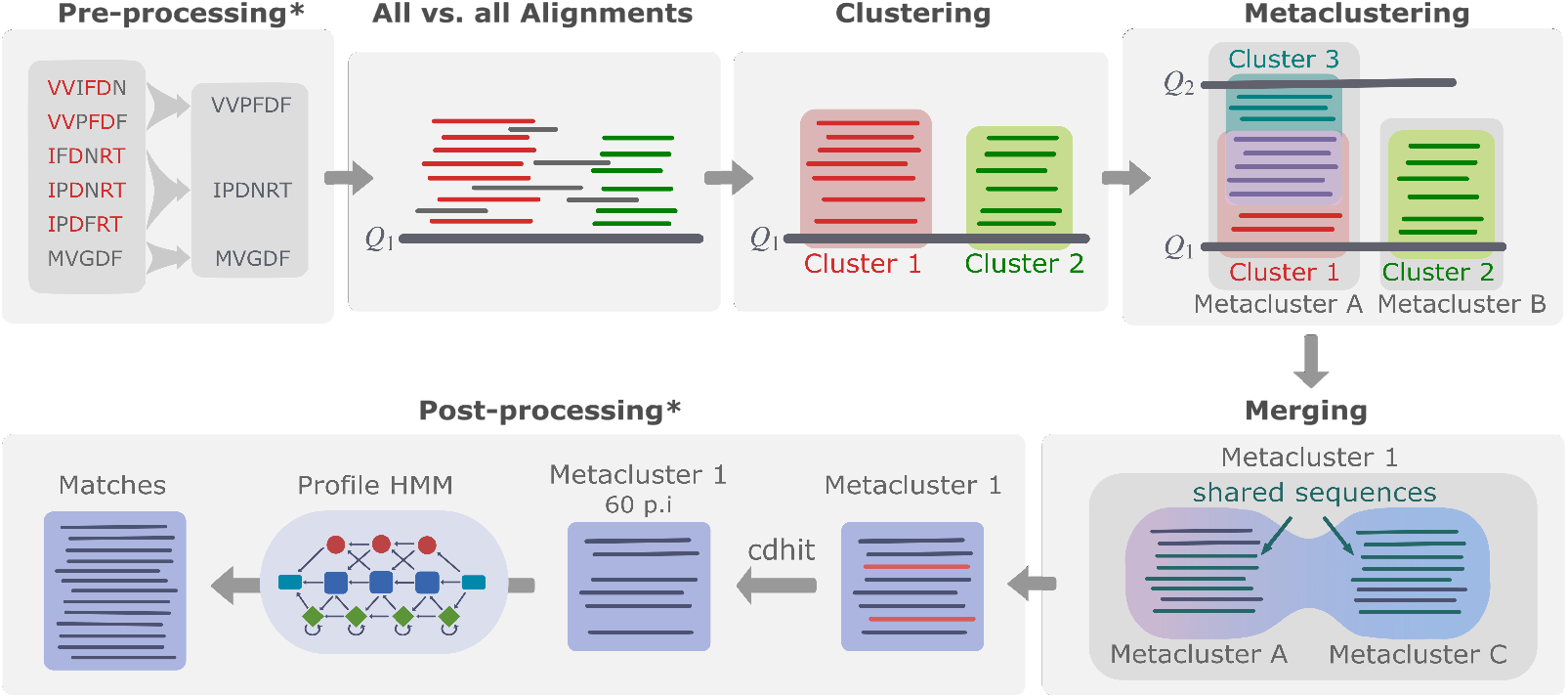
Steps of the DPCfam pipeline. **Pre-processing**: we take all sequences in UHGP-50 or the redundancy-reduced version of UHGP. **All-versus-all alignment**: for each sequence in UHGP-50 (query sequence), local alignments are generated against all other UHGP-50 sequences (search sequences) using blastp. **Primary clustering**: the sequence search regions (SSRs) that align to a query sequence are grouped together using DPC according to the relative position of their alignments along the query sequence. **Metaclustering**: the primary clusters from different query sequences are grouped together to form metaclusters using DPC according to how many SSRs they share. **Merging and filtering**.: metaclusters that after the previous step still share a significant number of search sequence regions are merged together to obtain the final list of metaclusters. **Post-processing**: profile Hidden Markov Models are generated for each metacluster and run against UHGP-50 in order to increase coverage of the automatic classification.

### Pre-processing

The DPCfam algorithm takes as input a dataset of non-redundant sequences with at most 50% amino acid identity among its entries. The filtering procedure by sequence identity avoids the over-representation of the number of pairwise alignments of a given protein region which could introduce artefacts in the final output. For this study, we used the clustered dataset UHGP-50 version 1.0 provided by MGnify and available for download at https://www.ebi.ac.uk/metagenomics/.

### All-versus-all alignment

The first step of the pipeline is an all-versus-all local pairwise sequence alignment computed with blastp, a software part of the BLAST+ v. 2.2.30 suite [4]. The result is a collection of search sequence regions (SSRs) that align with each query protein. This step is by far the most computationally demanding, despite being embarrassingly parallel, as each query alignment search is independent.

### Primary clustering

For each query sequence *Q*, the output of blastp consists of a collection of SSRs that align to different portions of *Q*. The primary clustering procedure groups these SSRs based on their overlap along the query sequence. To this end, we first define the following distance measure between two generic SSRs *i* and *j* of a query sequence *Q*,

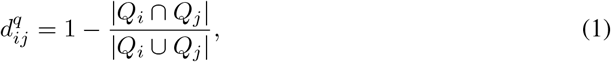

where *Q*_*i*_ and *Q*_*j*_ are the portions of the query *Q* covered by the respective SSRs. Note that if the SSRs cover exactly the same region of the query, the distance is zero, and if the two SSRs do not overlap, the distance is 1. Next, separately for each query *Q*, we use the above distance to cluster its SSRs using the DPC algorithm. Each cluster, or primary cluster (PC), represents a high-density region along the query sequence and comprises a set of SSRs. From a query perspective, primary clustering segments the sequence into regions akin to domains or domain combinations (architectures). This is also reflected on the associated SSRs, which represent domains or domain combinations of the search sequences they originate from. SSRs in primary clusters thus represent a first family-like group of sequences anchored to a specific query.

### Metaclustering (or Secondary Clustering)

Once we identify the segmentation of the query sequences according to clusters of SSRs, we proceed to classify them into metaclusters. We define the distance between two primary clusters as the number of SSRs they have in common with an overlap bigger than 0.8, normalized by the size of the smaller primary cluster. The secondary clustering procedure is done once again using the DPC algorithm, and the resulting metaclusters are collections of primary clusters which in turn contain a collection of SSRs.

### Merging and filtering

At this stage of the pipeline, we have a set of SSRs classified into metaclusters, each laying around a density peak of the landscape of primary clusters. Given the nature of the algorithm, two peaks separated by a high-density saddle point will be considered as independent metaclusters but could still share a large number of SSRs. To avoid this phenomenon we perform a merging procedure in which we join metaclusters that match this condition. We define the distance between two metaclusters as the average primary cluster distance of its members, defined in the previous section. If the average distance is smaller than 0.9 we merge the metaclusters. After the merging step we perform a filtering step to remove SSRs duplicates within MCs and, separately, to remove outliers. SSRs in metaclusters constitute the seeds of DPCfam-generated putative families.

### Post-processing

The output of the previous stage is a list of metaclusters, each containing a set of protein regions selected among the original SSRs. To further extend the sequence and residue coverage of our MCs, for each of them, we build a profile-HMM following procedure: 1) pruning of the MC seed sequences with CD-HIT v. 4.7 [13] at 60 percent identity; 2) generation from the pruned seed of a multiple sequence alignment (MSA) with the software MUSCLE v. 3.8.31 [23] from the pruned MCs; 3) construction from the MSA of a profile-HMM with HMMER - hmmbuild v.

3.1b2 [18]; 4) running of the profile-HMM against UHGP-50 using HMMER-hmmsearch (domain E-value=0.03, protein E-value=0.01). Note that the profile-HMM models can further be used to automatically annotate previously unseen proteins without the need to rerun the DPCfam clustering algorithm.

### Data Records

The full dataset is available for download from Zenodo (https://doi.org/10.5281/zenodo.7335147), and it can be explored interactively from our dedicated website at https://dpcfam.areasciencepark.it/uhgp. Only metaclusters with more than 50 protein seeds and an average sequence length of more than 50 amino acids are included in the dataset, resulting in a total of 10,778 metaclusters. To make the data more interoperable and reusable, metadata about all the seeds have been combined into a single XML file. This format allows for different types of information to be included in a hierarchical but flexible structure that can easily be built up sequentially. It is a machine-readable and well-established format used in many projects in the life sciences. In particular, it is the format used by the UniProt consortium to publish protein annotations. With this in mind, we also built an XML file containing all UHGP-50 proteins and annotated it with DPCfam-UHGP50 metaclusters and Pfam family membership.

In the remaining part of this section, we provide a detailed description of the content of the Zenodo repository, including the complete list of files associated with each metacluster and the XML files containing MCs information organized hierarchically along with UHGP-50 protein annotation.

### Metacluster Files

For each of the 10,778 metaclusters, we provide a fasta file with the metacluster seed sequences, a fasta file with the representative seed sequences after clustering at 60 percent identity, a fasta file with the multiple sequence alignment of the representative seed sequences, and a file with the corresponding profile-HMM. Files are grouped by category and compressed into the following archives: *seeds*.*zip, filtered_seeds*.*zip, metaclusters_msas*.*tar*.*gz*, and *metaclusters_hmms*.*tar*.*gz*.

### XML files

The *metaclusters_xml*.*tar*.*gz* archive contains the *metaclusters_uhgp50*.*xml* file and auxiliary parsing scripts, the use of which is described in *README*.*md*. In addition to the seed sequences, the XML file contains the following information about each metacluster: the number of seed sequences, the mean and standard deviation of the amino acid length of the seed sequences, the fraction of amino acids in a low-complexity region (LC), the fraction of amino acids in a coiled-coil region (CC), the fraction of amino acids in a disordered region (DIS) and the mean number of transmembrane regions (TM).

We also report data comparing the resulting UHGP-50 metaclusters with the Pfam annotation included in the UHGP-50 downloadable database. The metadata resulting from this comparison is described in Table 1. For an in-depth description of the analysis we refer to Technical Validation and Russo et al. [27]. Metaclusters are classified into four groups: equivalent, reduced, extended, and shifted. The specific category assigned to each metacluster depends on the fraction of the seed region that Pfam covers and vice versa. These values are averaged over all metacluster seed sequence - Pfam sequence pairs. If both are within 0.2, we consider the metacluster equivalent to the Pfam families it overlaps with; if both fractions are larger than 0.2, the metacluster is categorized as shifted. If only one of the fractions is larger than 0.2, the metacluster is either extended or reduced (see Figure 2).

**Table 1:** Detailed description for each XML field containing the DPCfam vs Pfam comparison metadata.

| XML Field Name | Description |
| --- | --- |
| DA | Dominant Pfam Architecture. Note: this field is UNK if no dominant architecture was found |
| DAC | Dominant Pfam Architecture (Clan level) |
| Percent_DA | Percent of sequences in the metacluster matching the dominant architecture |
| Percent_DAC | Percent of sequences in the metacluster matching the dominant architecture at a clan level |
| Percent_DACF | Percent of sequences in the metacluster matching the dominant architecture at a clan level, or a subset of it, including unannotated regions |
| Percent_DACFA | Percent of sequences in the metacluster matching the dominant architecture at a clan level, or a subset or superset |
| Label | A label to qualitatively describe the overlap type of the metacluster with the dominant architecture, which can be: equivalent, reduced, extended or shifted |
| Fred | Represents the fraction of the Pfam family that is not covered by the metacluster |
| Fext | Represents the fraction of the metacluster that is not covered by the Pfam family |
| Pfam_sequences | Number of sequences in the metacluster's seed used for comparison with Pfam. |

**Figure 2:**
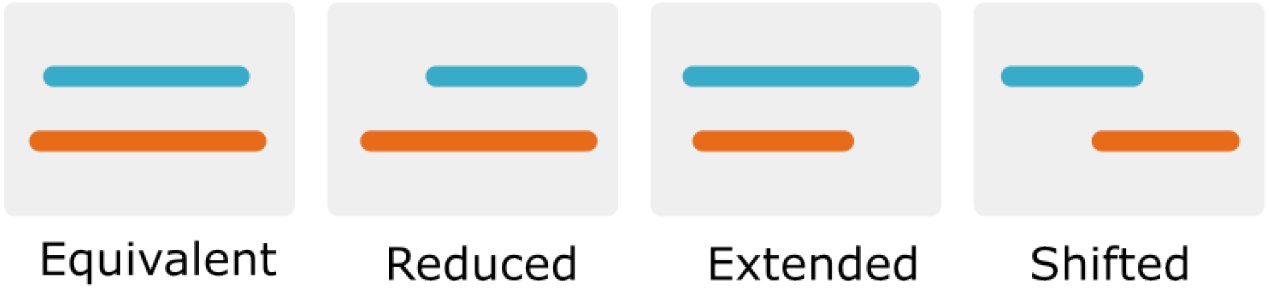
Visual representation of the different types of matches between a Pfam and DPCfam metacluster. The orange region represents the ground truth (Pfam), while the blue region represents the DPCfam metacluster. For a more detailed description of how these categories are assigned, refer to the original DPCfam article [27].

The archive *uhgp_xml*.*tar*.*gz* contains the UHGP-50 protein dataset annotated with Pfam and DPCfam families. For DPCfam, it is also indicated if the protein is a seed sequence or if it was found after scanning the database with the profile-HMMs.

### Technical Validation

First, we look into generic properties of the metaclusters, including size distribution, predicted average number of transmembrane regions, and the fraction of disordered, coiled-coil, and low-complexity regions. Second, using Pfam annotation as a reference, we test the existence of homologous relationships between MC members and analyse the quality of their boundaries. Finally, we study the overlap between UHGP-50 and UniRef50 metaclusters. From these analyses, we identified 1,261 unknown MCs, i.e. MCs that do not overlap with either Pfam or DPCfam-UniRef50 annotation. While overlapping metaclusters are helpful in improving existing annotation, the subset of unknown entries might represent protein families that are novel and potentially unique to the gut microbiome.

### Metaclusters general properties

Clustering UHGP-50 with DPCfam produced 40,738 metaclusters. The median MC size (i.e., number of SSRs) was 24, while the average size was 90.5. Figure 3 shows the Complementary Cumulative Distribution Function (CCDF) calculated from the metaclusters size distribution for UHGP-50 (blue) and, as a reference, UniRef50 (grey). The relationship between how frequent protein domain families appear in nature and their size was first described by Gerstein et al. [21] and can be explained by a birth, death, and innovation evolutionary model (BDIM) [11]. The resulting analytical expression is a Generalized Pareto Distribution which approximately fits the data with an exponent of 1.23*±*0.01 (red).

**Figure 3:**
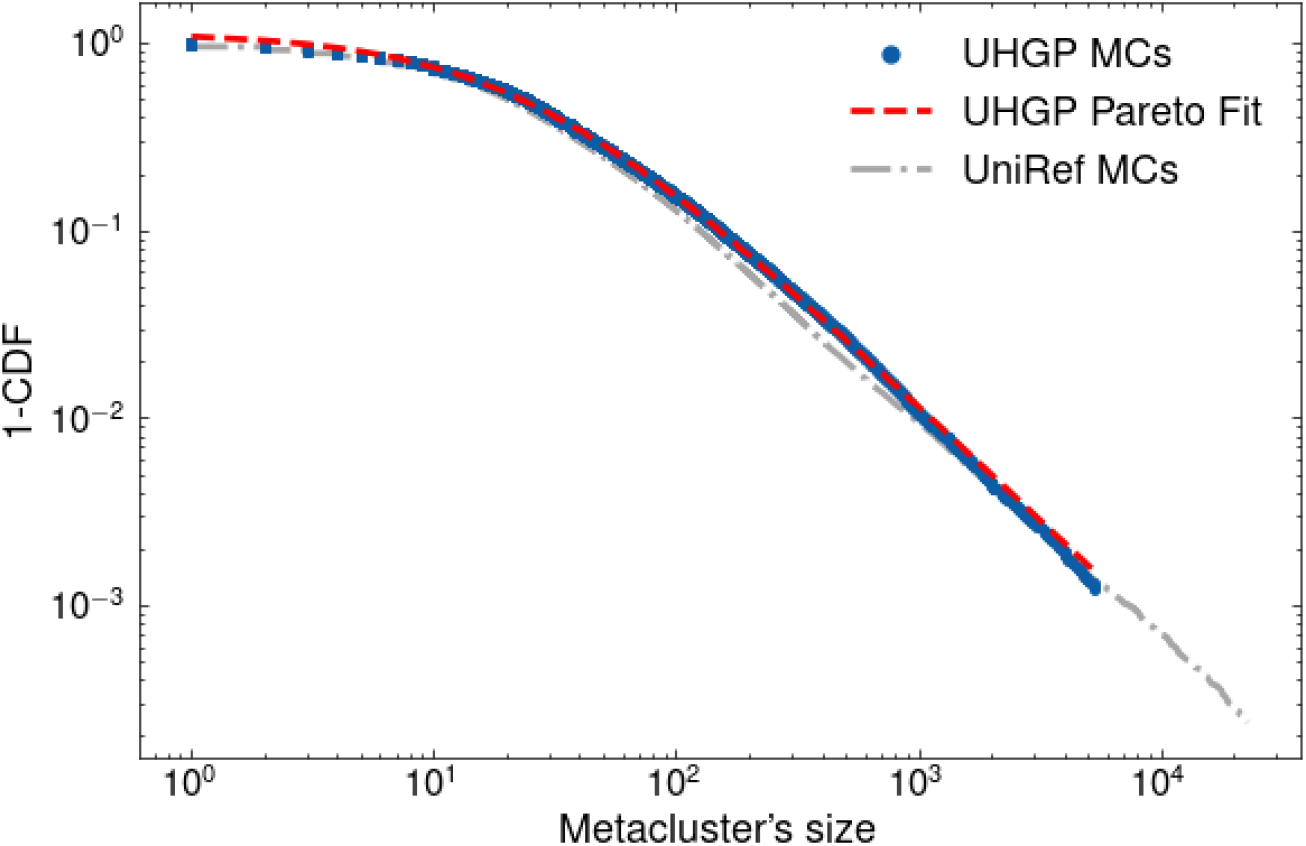
Log-Log plot of the Complementary Cumulative Distribution Function (CCDF) of UHGP-MCs (blue) and UR50-MCs (grey) sizes. The red line is the best fit of the data with a CCDF calculated from a Generalized Pareto Distribution (exponent of the Pareto 1.23*±*0.01).

In this regime of ‘the rich get richer’, elements larger in size are usually the more interesting ones. Moreover, as noted in our previous work [27], MCs of smaller size are, on average, of lower quality. Because of this, we decide to focus our downstream analysis on the set of MCs containing at least 50 members. We also excluded MCs with an average domain length of less than 50 amino acids since length also affects the quality of the alignments. After applying these filters, we obtained 10,778 metaclusters which, in the following, we refer to as the DPCfam-UHGP50 dataset. The median MC size after the filtering is 282.7, while the average size is 110.

Next, we examined the fraction of amino acids which are predicted to be found in a coiled-coil (CC), low-complexity (LC), and disordered (DIS) region, using the software DeepCoil[15], Segmasker[5] and IUPred2A[20] respectively. We labeled MCs with more than 10% of amino acids from all of their members in a low-complexity or coiled-coil region as Low Complexity MCs or Coiled Coil MCs, respectively. To label MCs as Disordered we used instead a threshold of 50% of all its amino acids in a disordered state. Figure 4 shows a Venn Diagram with the percentage of MCs in one or more of these categories. Although not a direct measure of quality, high values of these quantities indicate a bias in the amino acid composition of their member sequences, which makes it harder to infer homology from sequence similarity. This is especially true for low-complexity regions and coiled-coil regions.

**Figure 4:**
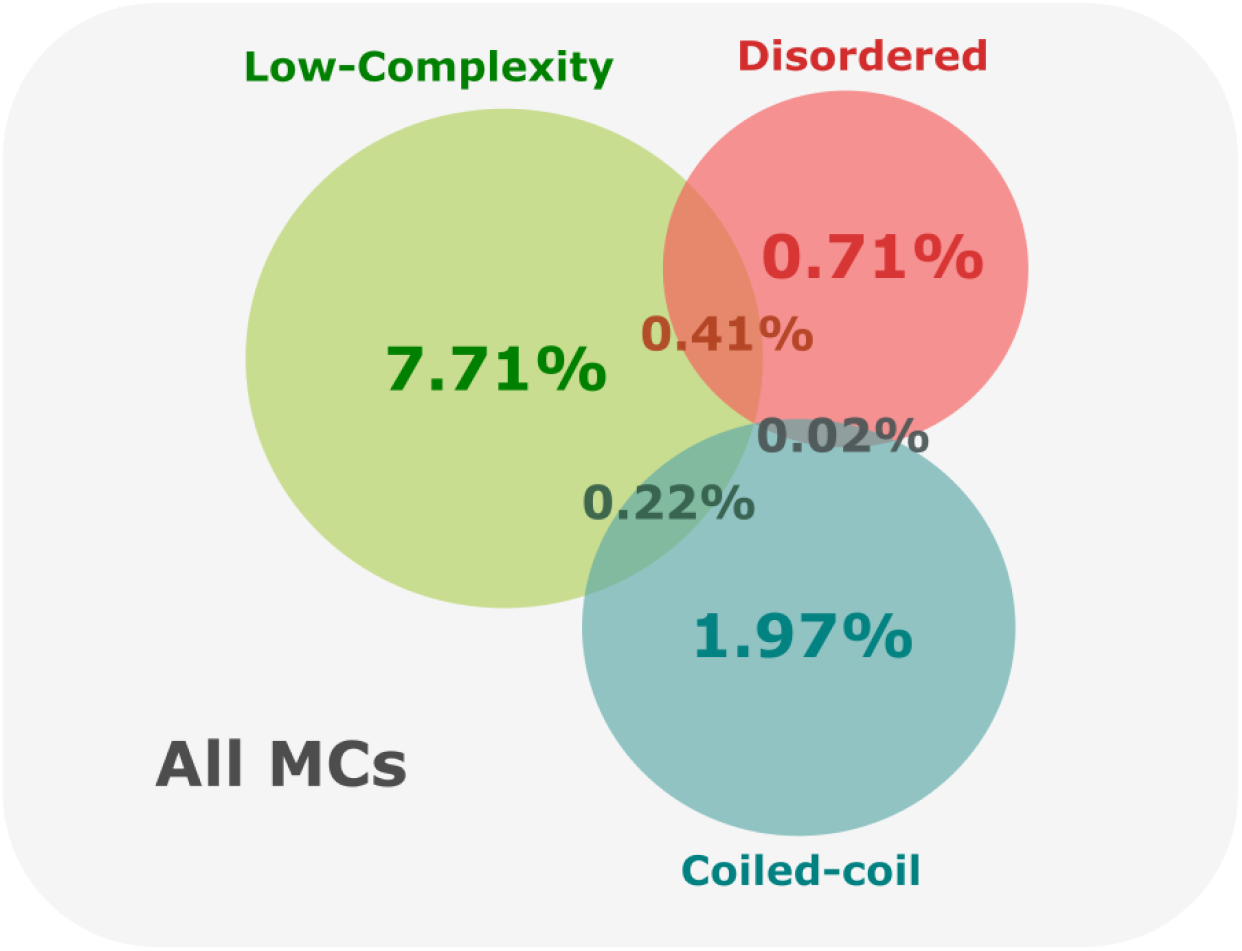
Percentage of metaclusters labeled as Low Complexity, Disordered, or Coiled Coil based on the fraction of amino acids of their seed sequences that are predicted to be found in a low-complexity, disordered, or coiled-coil region.

Results were in line with what we had observed when running DPCfam on UniRef50[27]; indeed, the majority of the MCs (88.96%) did not fall into any of these categories. We noted that the fraction of disordered MCs was rather small (about 1%). This was expected, however, since intrinsic disorder is not very common in prokaryotes[2], which constitute the vast majority of gut microbiome organisms[1]. As an additional measure, we used the software Phobius[12] to predict the number of transmembrane helical domains. 6.8% of MCs had an average of at least two transmembrane regions per domain and were thus likely to represent transmembrane helical bundle domains.

### Comparison with Pfam

To assess homologous relationships between metacluster seed sequences and to check how well our automatically-generated MCs boundaries mapped to those of manually annotated protein families, we compared our results with UHGP-50 Pfam annotation. Since DPCfam is a fully unsupervised method we do not expect complete agreement between both databases. However, for the cases where an overlap exists, it validates DPCfam results and its ability to identify protein families correctly.

To perform the comparison between the two classifications, we first extracted the Pfam labels from the Interpro scan file, which comes as part of the UHGP-50 downloadable database. In order to include multiple family architectures in the analysis, since MCs may extend over one or more families, we extended the Pfam annotation by combining single families into all possible family architectures. In what follows, we will use the term *Pfam architecture* to refer to single families or multiple family architectures.

To identify the Pfam architecture which best represents a particular DPCfam MC, we adopted the same criteria used in Russo et al. [27]. In particular, we defined the MC dominant architecture (DA) as the Pfam architecture shared by the largest number of its seed sequences (note that annotating a single amino acid is enough to associate a seed sequence to a Pfam family). To compare DPCfam and Pfam, we select only those MCs that have a DA shared by at least 50% of the seed sequences.

Figure 5 shows the distribution of the average overlap between each MC and its corresponding DA. The histogram includes the 4,434 metaclusters for which a Pfam DA*≥*50% was present. Colours correspond to the type of overlap between the two annotations (see Fig. 2). Among the MCs we consider here, 41.7% were classified as equivalent, 26.2% were reduced, 20.5% were extended and 11.6% were shifted.

**Figure 5:**
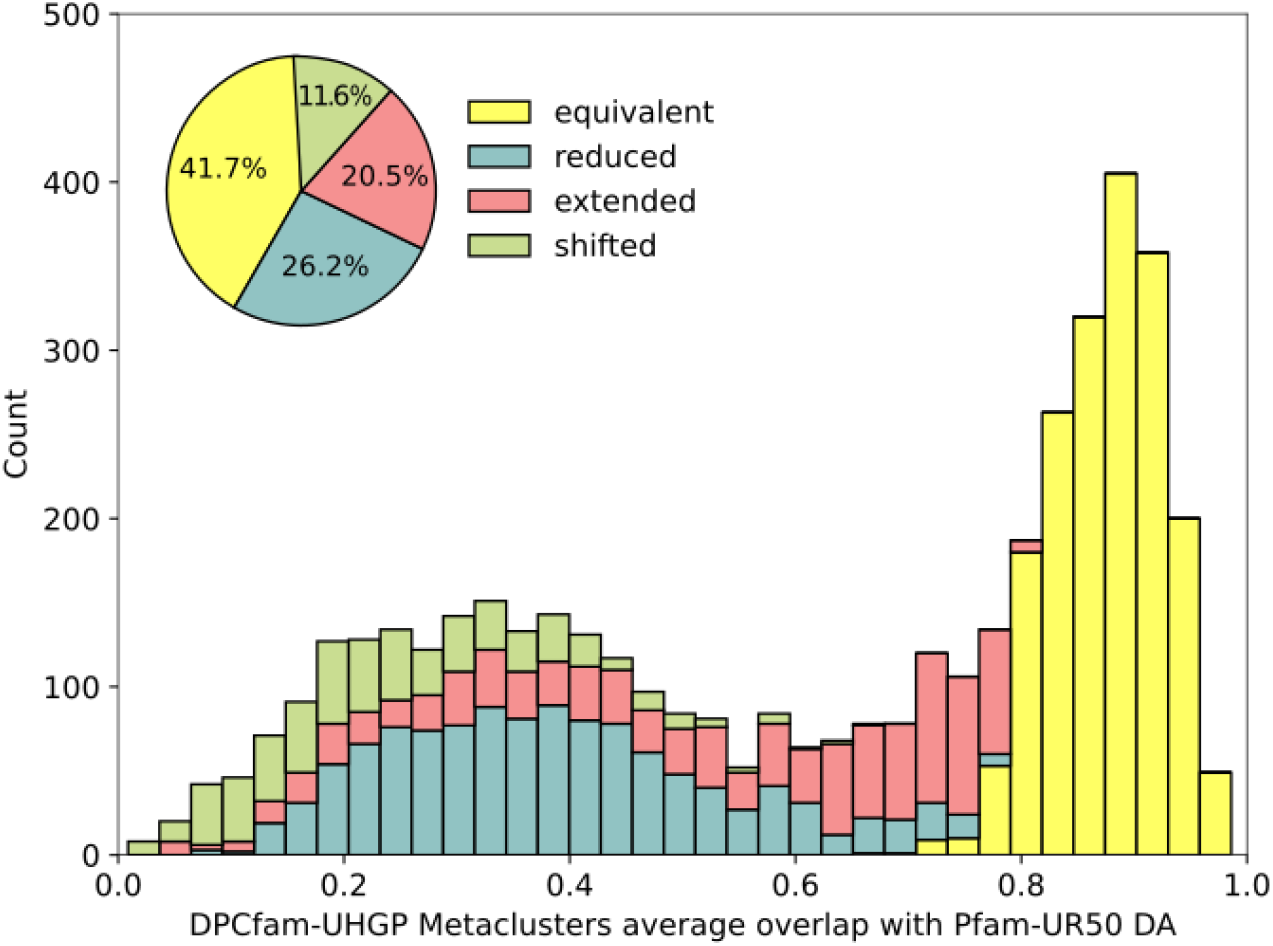
DPCfam-UHGP50 metaclusters average overlap with respect to its Pfam dominant architecture (only MCs with DA≥50%). Colors show the overlap class decomposition for each bin. The pie chart shows the aggregated contribution of each class expressed as a percentage of the 4,434 metaclusters.

Equivalent MCs have good boundary agreements with their Pfam DA. Reduced and extended metaclusters identify regions that are, on average, shorter or larger than the DA, respectively. Shifted metaclusters, conversely, correspond to cases where the MC covers regions that are consistently located N-terminal or C-terminal to the DA.

Among the remaining metaclusters without a well-defined DA, 2,277 do not overlap with any Pfam annotation and were therefore labeled as unknown to Pfam. This pool of MCs is particularly intriguing since it may include novel protein families.

### Comparison with DPCfam-UniRef50

To further evaluate the quality of our results, we compare DPCfam-UHGP50 metaclusters (UHGP-MC) against DPCfam-UniRef50 metaclusters (UR-MCs). The procedure was analogous to the one described in the previous section in which we compared UHGP-MCs against Pfam. The objective is to find, for each UHGP-MC, the correspondent UR-MC dominant architecture in order to establish a link between the two datasets. To obtain UR-MCs labels for UHGP-50 sequences, we ran the UR-MC profile-HMMs against UHGP-50 using HMMER - hmmsearch software.

Figure 6 shows the comparison results, where colours indicate the type of overlap with the DA. Among the 6,764 UHGP-MCs with a well-defined UR-MC dominant architecture, 50.6% are labeled as equivalent. This meant that most UHGP-50-generated MCs that had a well-matched UR-50-MC had close boundary correspondence with the latter. Only 8.7% of MCs were classified as shifted, further indicating that DPCfam can quite consistently identify family boundaries when run on different datasets.

**Figure 6:**
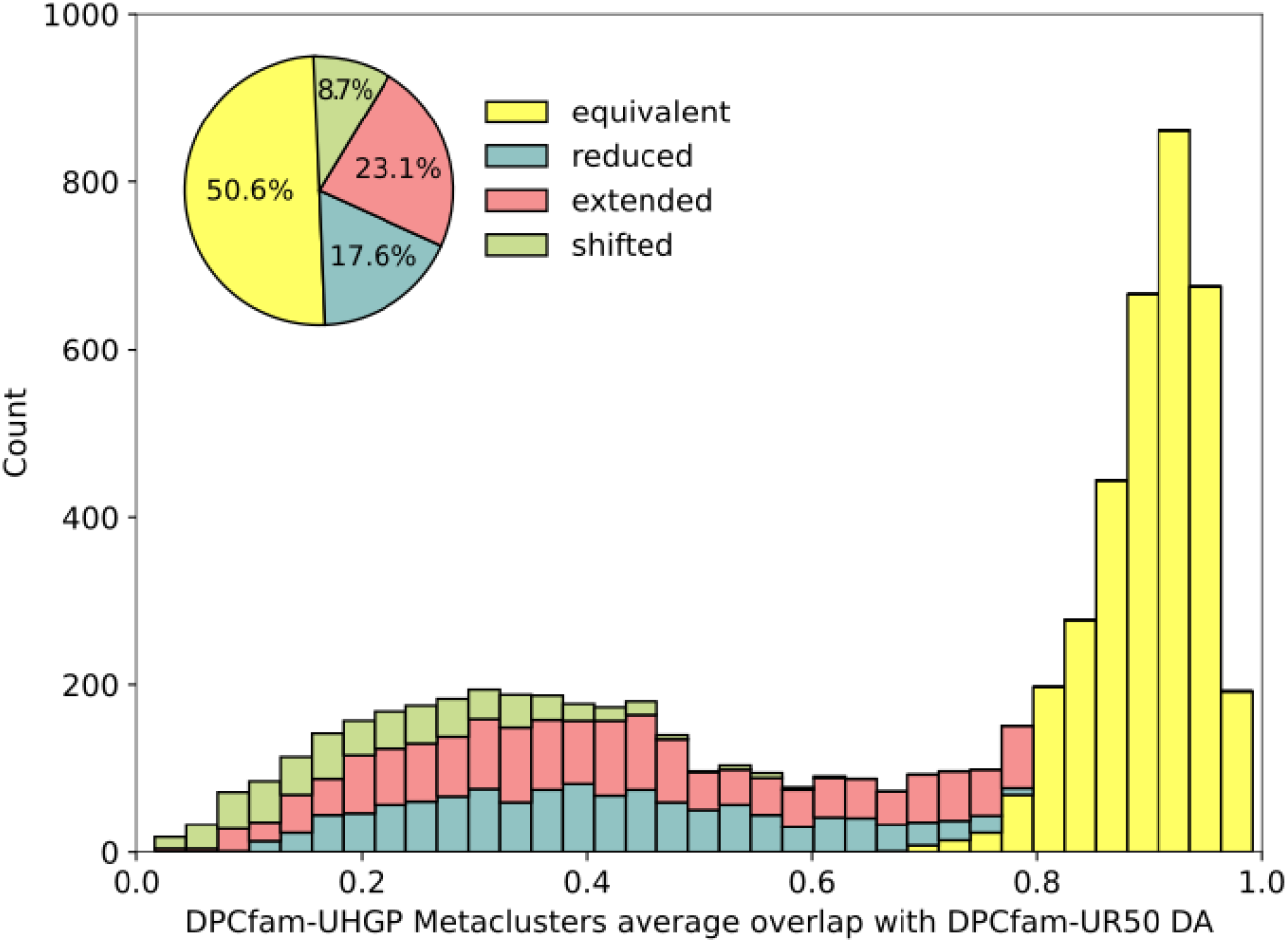
DPCfam-UHGP50 metaclusters average overlap with their DPCfam-UniRef50 dominant architecture (only MCs with DA*≥*50%). Colours show the overlap class decomposition for each bin. The pie chart shows the aggregated contribution of each class expressed as a percentage of the 6,764 metaclusters.

**Figure 7:**
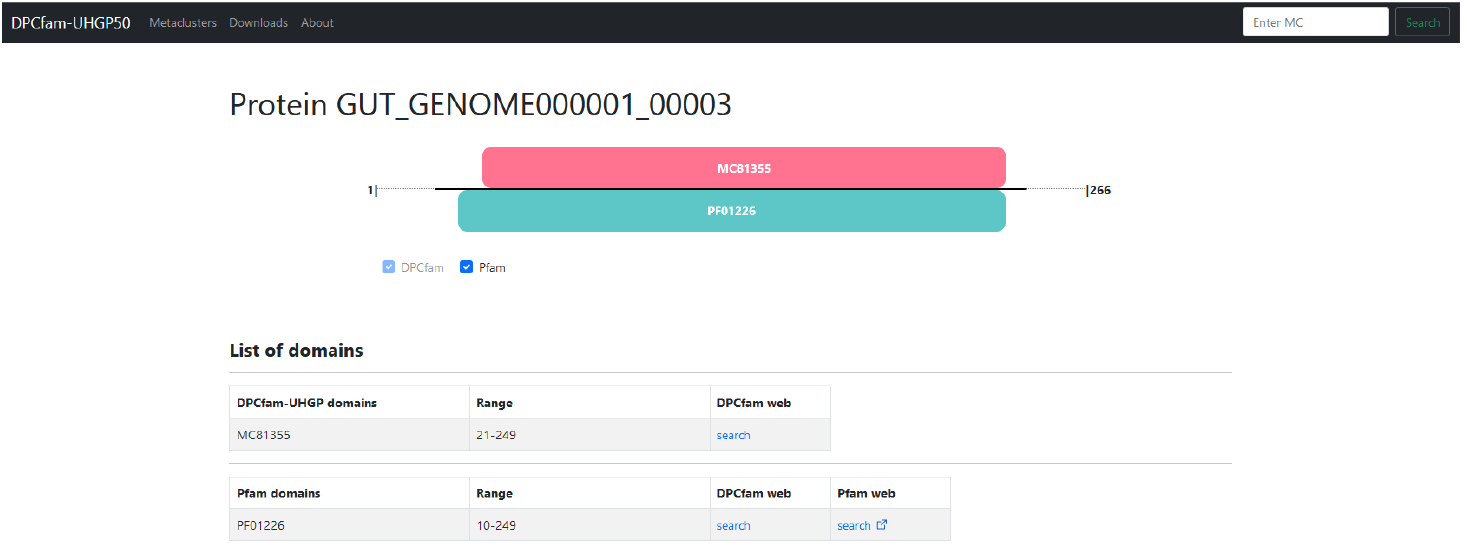
Screenshot of a protein entry page on the DPCfam web browser. A list of families is shown for the protein (both DPCfam and Pfam families) and their relative positions can be visualized in the schematic view.

Among those without a well-defined DA, we identified 1,953 with no overlapping UR-MC annotation. We labeled this subset of MCs as unknown to UR-MC. The overlap between the MCs unknown to both Pfam and UR-MC annotation creates a collection of potential new families that could be specific to the gut metagenome.

### Usage Notes

Parser scripts are included with the XML files to transform them into space-separated tables. We provide the information in XML format, which is better suited for the archival of different types of information in a single file; at the same time, we put the scripts at the user’s disposal for ease of use of the dataset, as tabular files can be more convenient to work with.

For the XML file containing the metaclusters, the parser script will generate a folder containing one fasta file per metacluster, as well as seven space-separated tables. Each fasta file contains all of the seed sequences of the corresponding metacluster. The space-separated txt files include the following: a list of all metacluster IDs, a table with statistical information on the seed sequences for each metacluster, a table with Pfam comparison measures for each metacluster, and four tables with additional metacluster properties (low-complexity, disordered, coiled-coil and transmembrane regions).

For the XML file listing all UHGP-50 proteins, the script generates a list of DPCfam matches (with start and end positions in the matching protein), a list of Pfam matches, a list of all UGHP-50 proteins, and lists of metacluster seed sequences before and after the CD-HIT filtering step.

Specific details on how to run the scripts are described in the README file. Parts of the code can be commented on to avoid generating all output files.

### Dedicated Webserver

The dataset can also be explored online on the dedicated DPCfam browser available at https://dpcfam.areasciencepark.it/uhgp. This website includes all the putative protein families from the DPCfam-UHGP50 clustering and lets the user search either for individual proteins or specific metaclusters, as well as Pfam families. The website also hosts information on previous DPCfam runs. Currently, it includes Pfam 33.0, DPCfam v1.1 on Uniref-50 and DPCfam v1.0 on UHGP-50.

Each DPCfam metacluster has its entry with information on basic statistics (including the percentage of low-complexity, coiled-coil, and disordered domains), the Pfam dominant architecture if present, as well as the complete list of seed sequences. The seed list, multiple sequence alignments, and HMM model can be downloaded from the metacluster page.

Proteins also have individual entries that summarize relevant information on them. For each protein, the corresponding entry page displays the list of DPCfam and Pfam domains and a graphical representation of their positions within the protein sequence. Results for Pfam queries include the matching DPCfam metaclusters if present and the degree and type of overlap.

## Code availability

The DPCfam pipeline source code is available for download at https://gitlab.com/area7/DPCfam/dpcfam. The repository includes usage notes and example datasets to test the algorithm.

## Acknowledgements

The authors acknowledge the AREA Science Park supercomputing platform ORFEO made available for conducting the research reported in this paper and the technical support of the Laboratory of Data Engineering staff. We thank Alessandro Laio for the conversations on the material of this paper. F.B., E.T.R. and E.N.V.G. were supported by the grant BOL “BIO Open Lab”. A.A. and A.C. were supported by the ARGO funding program.

## Notes

https://dpcfam.areasciencepark.it/uhgp

https://zenodo.org/record/7335147

### Competing Interest Statement

The authors have declared no competing interest.

